# *Aspergillus fumigatus* cytochrome c impacts conidial survival during sterilizing immunity

**DOI:** 10.1101/2023.06.07.544103

**Authors:** Matthew R James, Mariano A Aufiero, Elisa M Vesely, Sourabh Dhingra, Ko-Wei Liu, Tobias M Hohl, Robert A Cramer

**Author notes:** Address correspondence to Robert A Cramer and Tobias M Hohl, and. Present address: Sourabh Dhingra, Clemson University, Biological Sciences, Clemson, South Carolina, USA. Matthew R James and Mariano A Aufiero contributed equally to this work. Author order was determined by seniority on the project.

## Abstract

Invasive pulmonary aspergillosis (IPA) is a life-threatening infection caused by species in the ubiquitous fungal genus *Aspergillus*. While leukocyte-generated reactive oxygen species (ROS) are critical for the clearance of fungal conidia from the lung and resistance to IPA, the processes that govern ROS-dependent fungal cell death remain poorly defined. Using a flow cytometric approach that monitors two independent cell death markers, an endogenous histone H2A:mRFP nuclear integrity reporter and Sytox Blue cell impermeable (live/dead) stain, we observed that loss of *A. fumigatus* cytochrome c (*cycA*) results in reduced susceptibility to cell death from hydrogen peroxide (H_2_O_2_) treatment. Consistent with these observations *in vitro*, loss of *cycA* confers resistance to both NADPH-oxidase -dependent and -independent killing by host leukocytes. Fungal ROS resistance is partly mediated in part by Bir1, a homolog to survivin in humans, as Bir1 overexpression results in decreased ROS-induced conidial cell death and reduced killing by innate immune cells *in vivo*. We further report that overexpression of the Bir1 N-terminal BIR domain in *A. fumigatus* conidia results in altered expression of metabolic genes that functionally converge on mitochondrial function and cytochrome c (*cycA*) activity. Together, these studies demonstrate that *cycA* in *A. fumigatus* contributes to cell death responses that are induced by exogenous H_2_O_2_ and by host leukocytes.

**Importance:** *Aspergillus fumigatus* can cause a life-threatening infection known as invasive pulmonary aspergillosis (IPA), which is marked by fungus-attributable mortality rates of 20%-30%. Individuals at risk of IPA harbor genetic mutations or incur pharmacologic defects that impair myeloid cell numbers and/or function, exemplified by bone marrow transplant recipients, patients that receive corticosteroid therapy, or patients with Chronic Granulomatous Disease (CGD). However, treatments for *Aspergillus* infections remains limited, and resistance to the few existing drug classes is emerging. Recently, the World Health Organization (WHO) classified *A. fumigatus* as a critical priority fungal pathogen. Our research identifies an important aspect of fungal biology that impacts susceptibility to leukocyte killing. Furthering our understanding of mechanisms that mediate the outcome of fungal-leukocyte interactions will increase our understanding of both the underlying fungal biology governing cell death and innate immune evasion strategies utilized during mammalian infection pathogenesis. Consequently, our studies are a critical step toward leveraging these mechanisms for novel therapeutic advances.

## Introduction

*Aspergillus fumigatus* is a saprophytic mold that is ubiquitous in the environment. *A. fumigatus* can cause a life-threatening lung infection known as IPA (1). IPA manifests due to defects in innate immune cell numbers and function that results in the failure to clear inhaled fungal conidia prior to germination and hyphal tissue invasion in the lung (2–4). Notably, individuals with CGD have impaired phagocyte NADPH oxidase function, and neutrophils from these individuals mount a defective respiratory burst against various microbial pathogens (5, 6). The predisposition of individuals with CGD to IPA (and the susceptibility of CGD mice to *Aspergillus* challenge) indicate that products of NADPH oxidase are required for *A. fumigatus* clearance from the lung (7, 8). While it is known that host-generated reactive oxygen species (ROS) are critical for sterilizing immunity in the lung, and that conidia that undergo cell death from leukocytes display markers of regulated cell death (9), the mechanisms that mediate fungal cell death from host – generated ROS remain ill-defined.

Cytochrome c is a key regulator of apoptosis in Metazoa. Its release from the mitochondria under cell death inducing conditions, such as ROS exposure, promotes APAF-1-dependent activation of caspases (10, 11). Caspases are a family of cysteine proteases with specificity to aspartic acid residues that promote cell death via the controlled degradation of cellular proteins during apoptosis (12–14). Caspases are regulated by inhibitor of apoptosis proteins (IAPs) that directly inhibit caspase activity via a conserved Baculovirus inhibitor of apoptosis Repeat (BIR) domain (15–17). It was previously observed that loss of cytochrome C in *A. fumigatus* (encoded by the *cycA* gene) results in a strain capable of long-term persistence in the murine lung, resistance to murine macrophage mediated killing *in vitro,* and resistance to the superoxide generator menadione (18). Additionally, overexpression of Bir1, a homolog of the human IAP survivin, resulted in reduced susceptibility to both exogenous ROS and *in vivo* leukocyte killing (9). While *A. fumigatus* contains two genes canonically associated with caspase mediated cell death mechanisms in the Metazoa, fungi do not encode caspases and questions remain regarding the role of fungal cytochrome C and survivin homologs in fungal cell death mechanisms.

In this study, we further explore the impact of cytochrome C on *A. fumigatus* susceptibility to reactive oxygen species and host leukocytes. Using a flow cytometry approach that monitors two independent cell death markers over time, we observe that loss of *cycA* confers reduced susceptibility to hydrogen peroxide (H_2_O_2_) – induced conidial cell death. Furthermore, by using the **FL**uorescent **A**spergillus **RE**porter (FLARE) method (7), we observe that loss of *cycA* impacts infection in an immunocompetent murine infection model and confers reduced susceptibility to both NADPH oxidase-dependent and -independent killing by lung neutrophils. We further report that that overexpression of the Bir1 N-terminal BIR domain results in altered transcript levels of a subset of genes that converge on mitochondrial function and cytochrome c. These results further our understanding of host-leukocyte mediated fungal gene dependent cell death in the setting of sterilizing immunity and open new questions about the role of fungal mitochondrial integrity and/or function in these processes.

## Results

### *CycA* Amino Acid Sequence Annotation and Conservation

Cytochrome c is a highly conserved protein well known for its canonical function in the electron transport chain as an electron transporter between complex III and complex IV and as a mediator of metazoan apoptosis (10, 11, 19–21). Upon studying the *A. fumigatus* annotation of this gene in FungiDB, we noticed that the *cycA* amino acid sequence (Afu2g13110) was reported as 235aa in length, nearly double that of known cytochrome c proteins in eukaryotes (22). To resolve this unexpected finding, we obtained the AF293 reference nucleotide sequence from FungiDB and compared it to mRNA sequencing reads from Kowalski et al. 2019 (23). By analyzing sequencing reads in this data set we identified 4 predicted introns: intron 1 (G29 – G421), intron 2 (G454 – G513), intron 3 (G673 – G730), and intron 4 (G826 – G896). The revised amino acid sequence is 112 aa in length with a newly predicted TAA stop codon at T916 (Figure 1a). This new sequence displayed better sequence coverage to cytochrome c proteins in other model organisms (Figure 1b). When aligning the original *cycA* amino acid sequence, we observed only 41% coverage for *Aspergillus niger*, *Neurospora crassa*, *Arabidopsis thaliana*, and *Homo sapiens*. Using the revised amino acid sequence, we observed a much higher sequence similiarity of the *A. fumigatus* CycAp with other *Aspergillus* species orthologs; *A. niger* = 98%, *N. crassa* = 92%, *A. thaliana* = 89%, and *H. sapien* = 89%. Moreover, the revised amino acid sequence displayed high structural similarity to human *CycS* (Figure 1c).

**Figure 1.**
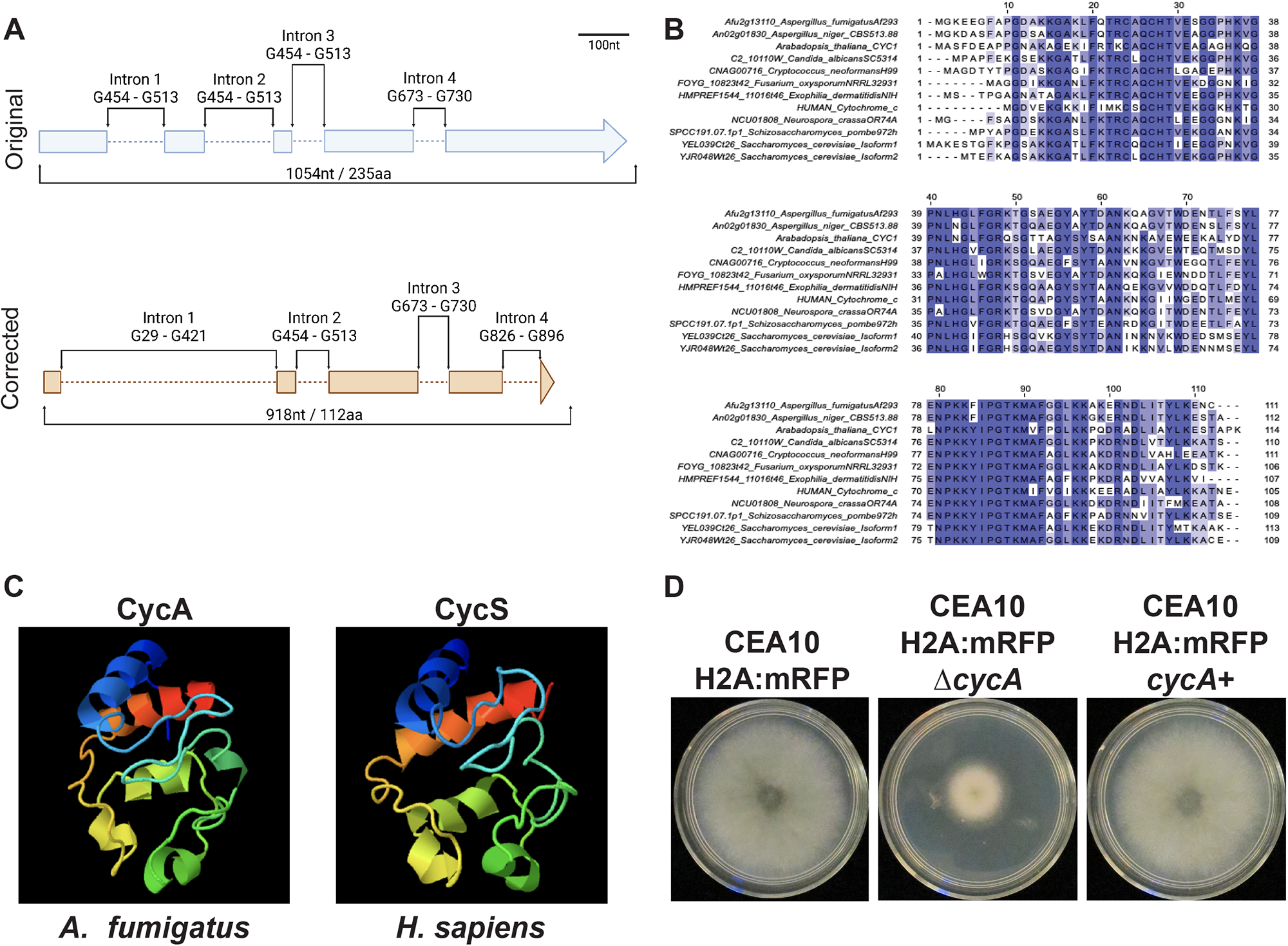
Cytochrome c sequence annotation in *A. fumigatus*. (A) Original and corrected annotation of cytochrome c in *A. fumigatus*. (B) Alignment of *A. fumigatus* cytochrome c to the sequences of other common organisms. (C) Phyre2 protein structure prediction of *A. fumigatus* cytochrome c protein (CycA) compared to human cytochrome c (CycS). (D) Representative images of the isogenic set used in this paper: CEA10 H2A:mRFP background strain, CEA10 H2A:mRFP Δ*cycA*, CEA10 H2A:mRFP Δ*cycA/cycA+*.

### Loss of *cycA* confers resistance to exogenous H_2_O_2_ treatment and alters cell death markers

Previous work in our lab identified a role for *cycA* in the oxidative stress response, whereby loss of the *cycA* gene conferred resistance to the superoxide generator menadione (18). Since overexpression of full-length or of the N-terminal domain of Bir1 conferred resistance to hydrogen peroxide (H_2_O_2_) in swollen conidia (9) we tested the hypothesis that loss of *cycA* also confers resistance to H_2_O_2_ in swollen conidia. To analyze conidial cell death, we first generated a histone H2A:mRFP nuclear reporter strain that uses loss of mRFP fluorescence as a marker of conidia cell death (CEA10 H2A:mRFP) (9, 24, 25) (Figure 2b). Using this strain, we generated the *cycA* null mutant CEA10 H2A:mRFP Δ*cycA* (referred to in this work as Δ*cycA*) by replacing the entire *cycA* gene with a hygromycin resistance cassette. We subsequently generated the reconstituted strain (*cycA*^+^) in the Δ*cycA* background. Consistent with previous work (18), we observed a marked growth defect in the Δ*cycA* strain as compared to the CEA10 H2A:mRFP background strain that was reconstituted by reinsertion of the *cycA* gene (Figure 1d).

**Figure 2.**
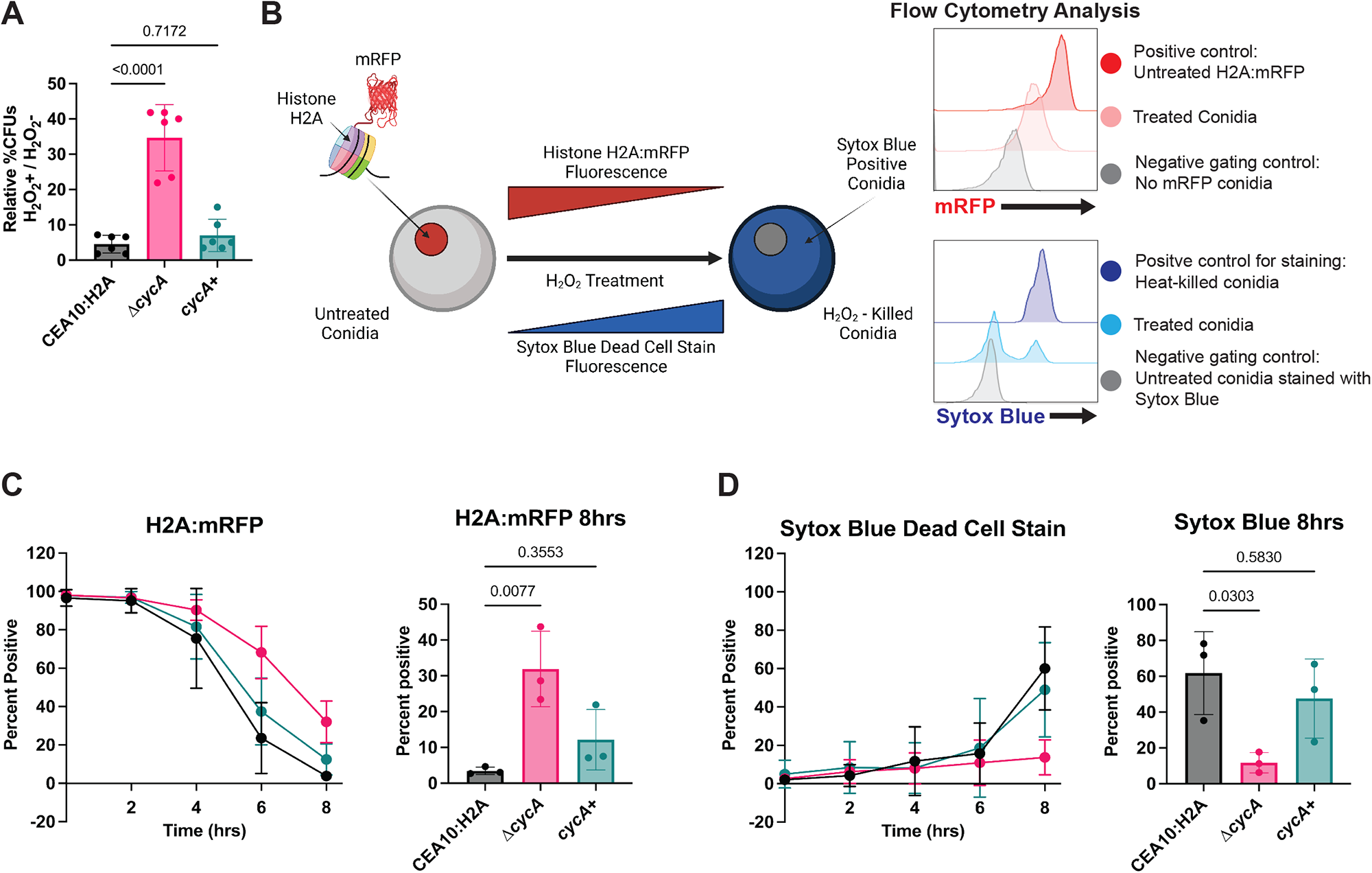
Loss of *cycA* alters multiple cell death markers after treatment with H_2_O_2_. (A) Normalized percent CFU’s. Percent untreated = (5mM treated CFUs / untreated CFUs). Data is representative of two biological repetitions including 3 technical repetitions each. Statistical analysis: one way ANOVA with Bartlett’s test. (B) Schematic of flow cytometry – based cell death assay experimental design. Over 8hrs treatment with 10mM H_2_O_2_, H2A:mRFP conidia were analyzed by flow cytometry for mRFP florescence and sytox blue florescence by flow cytometry. (C) Percent of histone H2A:mRFP positive conidia through H_2_O_2_ treatment (left) and quantification of histone H2A:mRFP positive cells at 8hrs (right). (D) Percent of Sytox Blue positive conidia through H_2_O_2_ treatment (left) and quantification of sytox slue positive cells at 8hrs (right). Data for C and D is representative of three biological repetitions including 15000 events each. Statistical analysis: one way ANOVA with Bartlett’s test.

Using this isogenic set, we observed that loss of *cycA* conferred an increase in viability after treatment with H_2_O_2_ as determined by the formation of CFUs (Figure 2a). We next monitored two independent markers of conidia cell death over time; the endogenous H2A:mRFP signal and Sytox blue dead cell stain (26) (Figure 2b). Every two hours, we stained a sample of H_2_O_2_ – treated conidia, and analyzed these for H2A:mRFP and Sytox blue fluorescence via flow cytometry. In the CEA10 H2A:mRFP strain, we observed that the majority of H2A:mRFP fluorescence signal loss occurred between 4 hrs and 6 hrs, consistent with previous observations (9). Between 6 hrs and 8 hrs we observed a significant increase in Sytox blue staining. For the Δ*cycA* strain, we observed a relative RFP fluorescence signal preservation during the treatment window, as judged by the presence of 31.9% mRFP^+^ cells compared to 3.48% (p = 0.0077) mRFP^+^ cells in the CEA10 H2A:mRFP background strain (Figure 2c). In addition to preserved H2A:mRFP fluorescence, we observed that the Δ*cycA* strain had only 11.67% Sytox blue^+^ conidia, while the CEA10 H2A:mRFP displayed 61.77% Sytox blue^+^ conidia (p = 0.0303) (Figure 2d). Taken together, these data suggest that loss of *cycA* reduces conidial susceptibility to H_2_O_2_ – induced cell death.

### Loss of *cycA* confers resistance to conidial killing by host phagocytes

NADPH oxidase-derived ROS are essential for host defense to *A. fumigatus* (6). Consequently, we hypothesized that loss of *cycA* would render *A. fumigatus* resistant to phagocyte killing in the lung. We tested this hypothesis by challenging C57BL/6J mice with either 3x10^7^ WT, *ΔcycA*, or *ΔcycA+* conidia and analyzing lung phagocyte numbers and phagocyte uptake and killing of conidia at 36 hours post fungal challenge, using a fungal bioreporter, termed FLARE, as previously described (7). FLARE conidia encode RFP as a sensor of fungal cell viability and are labeled with Alexa Fluor 633 (AF633) as a tracer fluorophore. Thus, RFP^+^AF633^+^ leukocytes contain live fungal cells, while RFP^-^AF633^+^ leukocytes contain killed fungal cells. With this method, all AF633^+^ leukocytes represent leukocytes with phagocytosed fungal cells, irrespective of RFP fluorescence. The frequency of leukocytes with live fungal cells (AF633^+^RFP^+^) divided by the frequency of fungus-engaged leukocytes (all AF633^+^ leukocytes; fungus-engaged leukocytes) represents a measure of fungal cell killing.

Compared to the *ΔcycA+* strain, there were fewer neutrophils in the lungs of mice challenged with the *ΔcycA* strain, but there was no difference in lung monocytes or monocyte-derived dendritic cells (Mo-DCs) (Supplementary Figure 1). We observed a slight decrease in neutrophil conidial uptake in mice challenged with the *ΔcycA* strain. However, phagocytosis by monocytes or Mo-DCs was not affected by loss of *cycA* (Figure 3a and 3b). Strikingly, the fraction of fungus-engaged neutrophils and monocytes that contained live conidia was dramatically increased in mice challenged with the *ΔcycA* strain compared to WT or with *ΔcycA+* (Figure 3a and 3c). This result suggests that loss of *cycA* confers resistance to killing by lung myeloid phagocyte cells, which are essential for anti-*Aspergillus* host defense.

**Figure 3.**
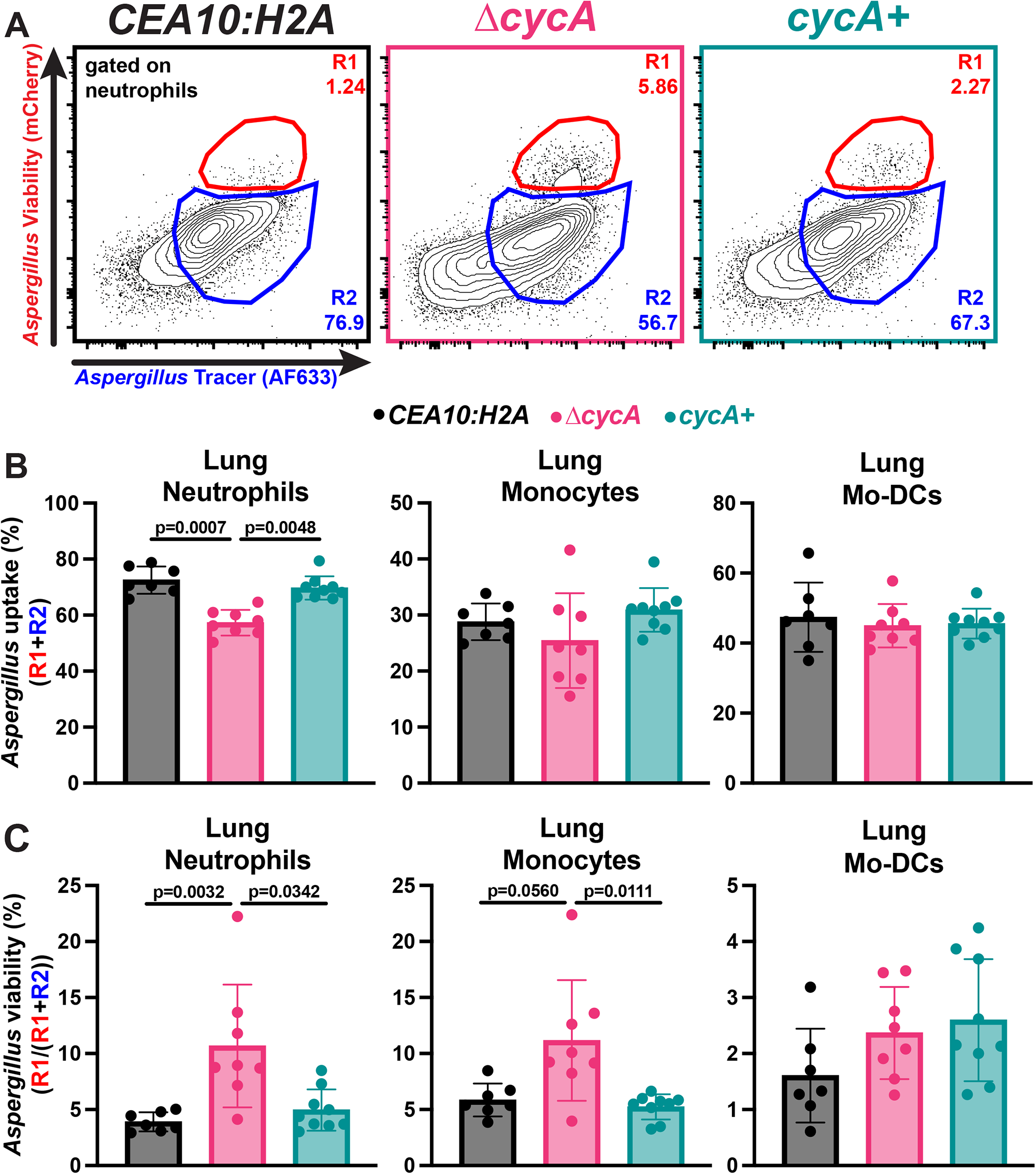
ΔcycA *A. fumigatus* is resistant to lung leukocyte killing. (A) Representative flow plots of lung neutrophil conidial uptake and killing, analyzed based on RFP and AF633 fluorescence, 36 hpi with 3×10^7^ AF633+ *CEA10:H2A* (black), *ΔcycA* (magenta), or *ΔcycA+* (teal) conidia. (B and C) The scatter plots indicate (B) conidial uptake (R1+R2) by and (C) conidial viability (R1/(R1+R2)) in indicated leukocyte subsets. (B and C) Dots represent individual mice and data are expressed as mean ± SEM. Statistical analysis: Kruskal-Wallis test with Dunn’s multiple comparisons test.

To test whether loss of *cycA* confers resistance to host NADPH oxidase-dependent killing, we generated bone marrow chimeric mice with equal ratios of p91phox^+/+^ (an essential subunit of the NADPH oxidase complex^1^) and p91phox^−/−^ cells, and challenged these mice with WT, *ΔcycA*, or *ΔcycA+* strains (Figure 4a). As expected, the fraction of viable WT and *ΔcycA+* conidia increased in p91phox^-/-^ neutrophils compared to p91phox^+/+^ neutrophils, consistent with the known role of NADPH oxidase as a vital host mechanism for killing of *A. fumigatus* conidia in the lung (Figure 4b and 4c). Surprisingly, the percentage of viable *ΔcycA* conidia was also higher in p91phox^-/-^ neutrophils compared to p91phox^+/+^ neutrophils (Figure 4b and 4c), indicating that NADPH oxidase-dependent mechanisms do not exclusively drive resistance to neutrophil killing conferred by loss of *ΔcycA*. Our *in vitro* findings, which demonstrate enhanced resistance of *ΔcycA* to chemically induced oxidative stress, combined with these in vivo results, suggest that the loss of *cycA* enables resistance to both oxidative and non-oxidative host killing mechanisms.

**Figure 4.**
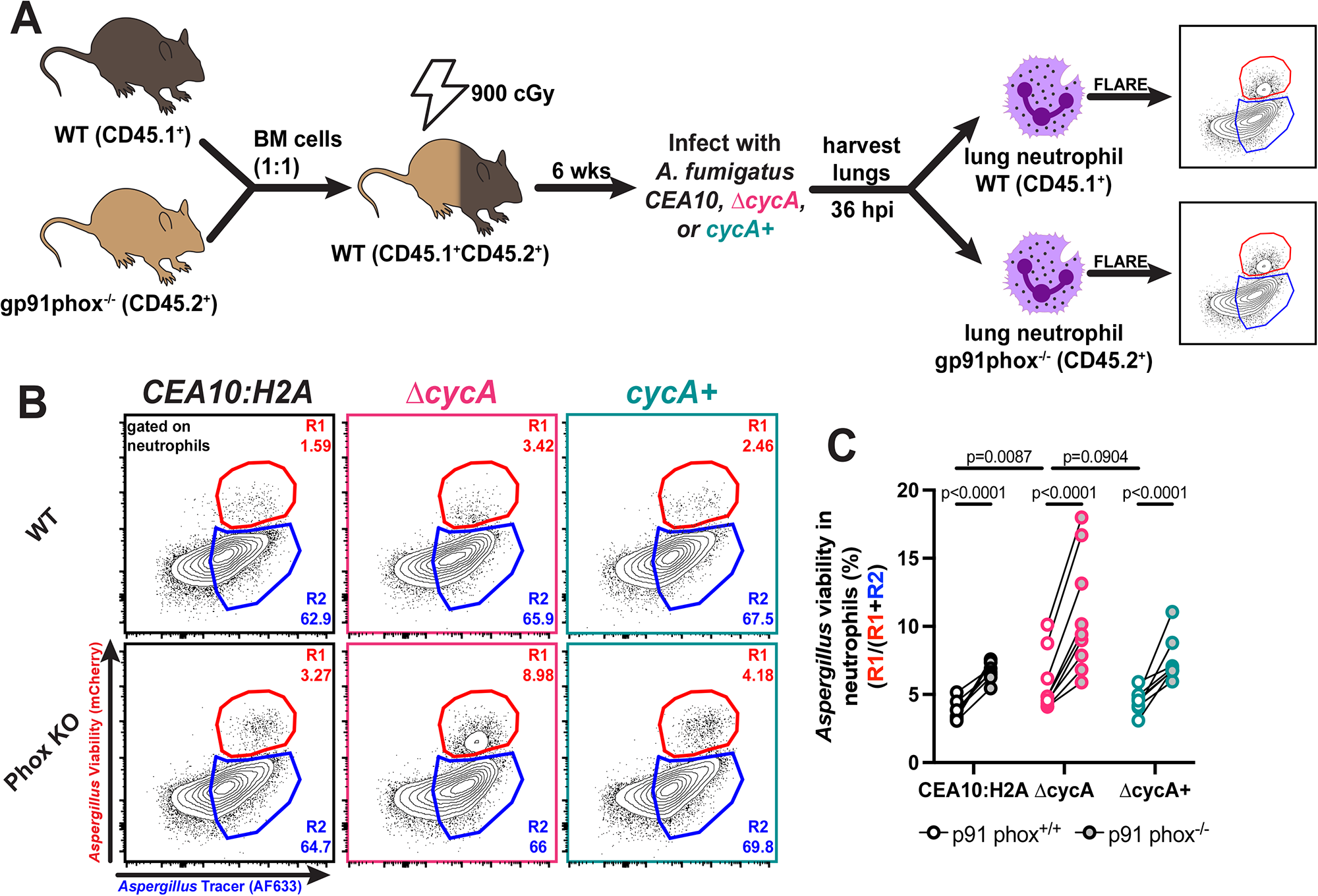
*ΔcycA* conidia viability in NADPH oxidase deficient leukocytes. (A) Mice received a 1:1 mixture of p91phox^+/+^ and p91phox^-/-^ bone marrow, were rested for 6 weeks, and then challenged with 3x10^7^ AF633-labeled *CEA10:H2A* (black), *ΔcycA* (magenta), or *ΔcycA+* (teal). (B) Representative flow plots of lung neutrophil conidial uptake and killing from mixed bone marrow chimera mice, analyzed based on RFP and AF633 fluorescence, 36 hpi with 3×10^7^ AF633+ *CEA10:H2A* (black), *ΔcycA* (magenta), or *ΔcycA+* (teal) conidia. (C) The scatter plot indicates conidial viability in p91phox^+/+^ (white center) and p91phox^−/−^ (gray center) neutrophils. Dots represent individual mice with mean ± SEM indicated. Statistical analysis: Blue lines indicate Wilcoxon matched pairs test; black lines indicate Kruskal-Wallis test with Dunn’s multiple comparisons test.

### RNA sequencing of Bir1^OE-N^ cells shows altered mRNA transcript level differences that suggests altered metabolic function

Recently, overexpression of the full-length (Bir1^OE^) or N-terminal Bir1 fragment that contains both BIR domains (Bir1^OE-N^) was observed to decrease *A. fumigatus* conidia susceptibility to exogenous H_2_O_2_ stress and result in increased fungal viability during leukocyte-fungal interactions in an immunocompetent murine infection model (9). These phenotypes are similar to the loss of *cycA*, and we next sought to more fully understand the genetic networks that dictate efficient oxidative stress induced killing of swollen fungal conidia. Therefore, we measured mRNA levels in Bir1^OE-N^ cells in comparison to the wild-type strain (ATCC46645-H2A:mRFP) prior to treatment with exogenous ROS (Figure 5a). RNA sequencing results indicated that transcripts of 413 genes were significantly increased, while 377 gene transcripts were decreased in mRNA abundance compared to WT cells (2-fold difference cutoff) (Figure 5b). mRNA abundance of genes involved in ‘Ribosome biosynthesis’ and ‘Mitochondrial activity’ in Bir1^OE-N^ cells were elevated compared to the wild-type strain by Gene Ontology (GO) Term analysis (Figure 5c). The gene with the largest fold increase in mRNA levels in Bir1^OE-N^ as compared to WT was *acoA*, which encodes for aconitate hydratase in *A. fumigatus* (<u>AFUB_001810</u>, increased 26.2-fold, indicated in Figure 5c). Moreover, we observed increased mRNA levels of genes that encode other key components of the electron transport chain in Bir1^OE-N^. These included complex II succinate dehydrogenase (<u>AFUB_041300</u>, 2.16-fold increased), complex III mitochondrial cytochrome b-c1 (<u>AFUB_058230</u>, 6.5-fold increased), mitochondrial cytochrome b-c1 (<u>AFUB_002450</u>, increased 6.8-fold), NADH-quinone oxidoreductase (<u>AFUB_006990</u>, 2.7-fold increased), NADH-ubiquinone oxidoreductase (<u>AFUB_068090</u>, 2.46-fold increased), and fumarate reductase (<u>AFUB_082020</u>, 4.5-fold increased) (Figure 5d). There data suggest alterations in electron transport function in Bir1^OE-N^ cells.

**Figure 5.**
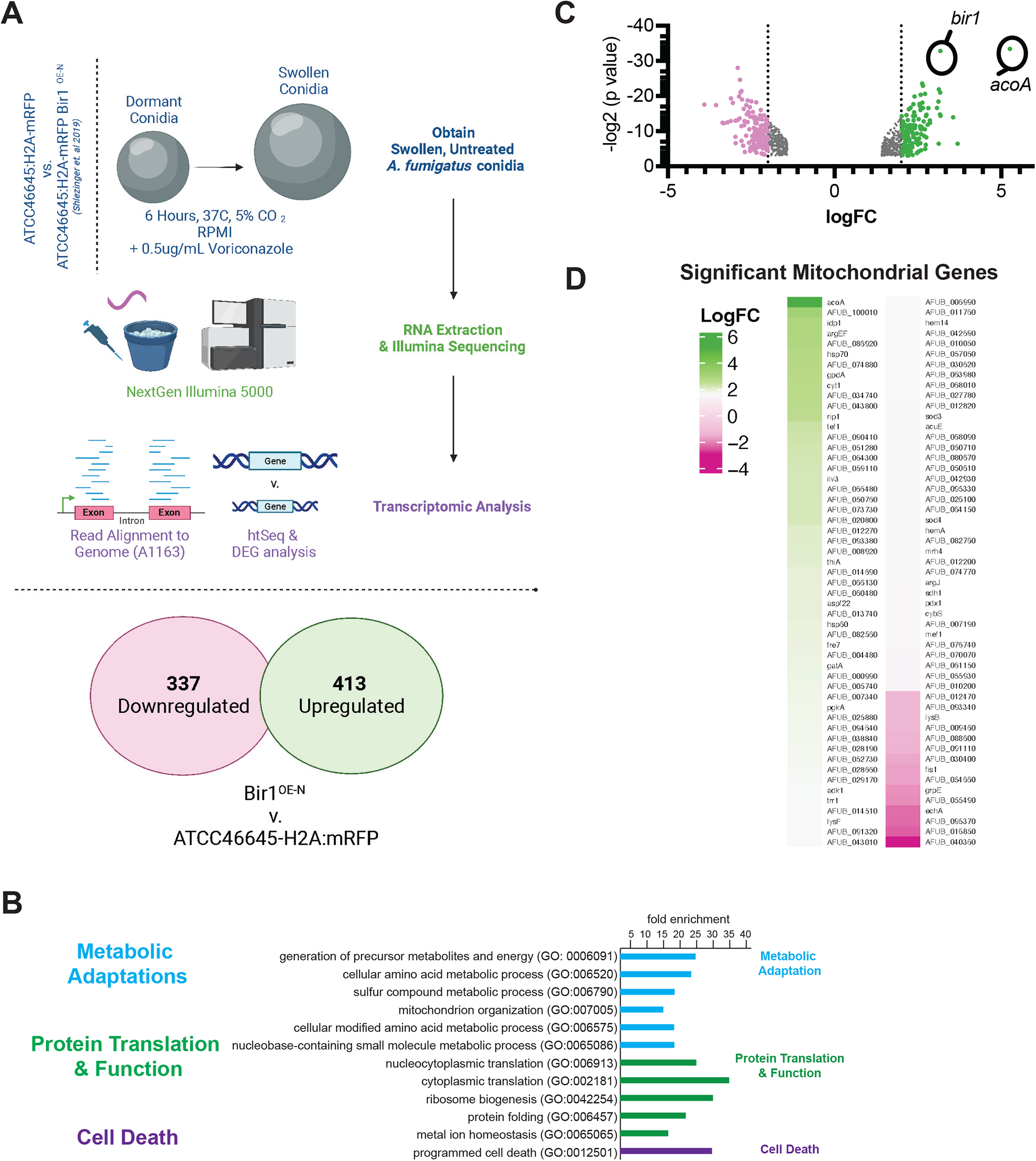
Oxidative stress-resistant Bir1^OE-N^ conidia show baseline transcriptional variation. Transcriptional profiling of swollen conidia of strain ATCC46645-H2A:mRFP compared with an isogenic strain that features artificially expressed AfBir1(Bir1^OE-N^) gene product (Shlezinger et al 2019). (A) Experimental set-up to determine gene expression in swollen conidia from both strains. (B) Differential gene expression comparing Bir1^OE-N^ to ATCC46645H2AmRFP, pink and green highlighting indicates genes downregulated and upregulated by more than logFC=2 respectively. (C) Using FungiDB GO Term analysis, GO ‘Mitochondria’ genes were compared to Bir1^OE-N^ DEGs and visualized by heat map.

We next examined mitochondrial genes with reduced mRNA levels in Bir1^OE-N^ vs. wild-type swollen conidia, and observed reductions in *echA* that encodes an Enoyl CoA-hydratase. A reduction in *echA* levels is consistent with a hypothesis that beta-oxidation may be suppressed in the utilized experimental conditions and that cells are not experiencing a ‘low glucose’ state before RNA extraction (27). In addition, we observed a significant reduction in transcript levels of the gene responsible for regulation of mitochondrial fission*, fis1* (AFUB_028960, reduced 3-fold), suggesting that potential alterations in mitochondrial fusion and fission dynamics in Bir1N-OE cells may occur. In immortalized human cells, reductions in mitochondrial fission have been linked to both decreased apoptotic cell death and decreased leakage of cytochrome c from the mitochondria (28, 29). Interestingly, genes for secondary metabolite biosynthesis showed a reduced mRNA transcript abundance (ex: polyketide F-9775B biosynthesis gene AFUB_033110, reduced 15.25-fold), which may indicate that Bir1^OE-N^ cells have a reduced biosynthesis of secondary metabolites under the conditions tested. Together, these data suggest that the leukocyte mediated cell death resistant Bir1^OE-N^ strain has altered transcriptional regulation of metabolic genes that functionally converge on cytochrome c activity and/or mitochondrial function.

## Discussion

Historically, research on regulated cell death (RCD) mechanisms has focused on metazoa (30–32), while RCD mechanisms outside of metazoa remain incompletely defined (33–35). In the fungi, a complicating factor is the lack of many canonical cell death regulators described in metazoans, such as caspases, caspase-interacting domains, and Bcl-2 family proteins, in fungal genomes. However, fungi encode homologs of mitochondrial-localized pro-apoptotic factors such as AifA, EndoG, HtrA2, cytochrome c, and homologs of gasdermin D and survivin (9, 25, 36–39). Aside from mammalian homologs, recent research suggests that cell death pathways that are unique to fungi exist. In *N. crassa*, the NLR-like protein PLP-1 functions in allorecognition by detecting SEC-9 from incompatible strains during heterokaryon incompatibility in germlings (40). In *N. crassa*, the gasdermin-like protein, RCD-1, can promote cell death during heterokaryon incompatibility the via interaction of two different alleles, *rcd-1-1* and *rcd-1-2*, which promotes the formation of pores on the plasma membrane (41). In *Saccharomyces cerevisiae*, AP-3 vesicle trafficking was implicated in cell death induced by acute heat stress, likely through function of the kinase Yck3 which leads to permeabilization of the vacuole (42). These results suggests that like metazoa, fungi may execute cell death processes in a regulated manner, though the mechanisms still remain to be fully determined.

To this end, studies in both plant and human fungal pathogens have suggested that alteration of Bir1 transcript levels impacts susceptibility to cell death and virulence (9, 25). In the necrotrophic fungal plant pathogen, *Botrytis cinerea*, overexpression of Bir1, or its N-terminal domain, conferred reduced susceptibility to cell death induced by plant defenses and resulted in increased infection lesion size on *Arabidopsis thaliana* or *Phaselus vulgaris* leaves (25). In the human fungal pathogen, *A. fumigatus,* overexpression of Bir1, or its N-terminal domain reduced H2A:mRFP loss and TUNEL staining, and increased viability when swollen conidia were exposed to exogenous H_2_O_2_. Moreover, overexpression of Bir1 was protective against sterilizing immunity from host leukocytes and increased virulence in a murine model of fungal bronchopneumonia (9). However, the mechanism by which increased Bir1 levels are protective against cell death in fungi remains to be defined. In fungi, it is unclear if Bir1 is directly involved in a regulated cell death pathway and/or involved the in regulation of cell processes and/or genetic networks that confer general oxidative stress resistance to fungal cells.

Previously it was reported that loss of cytochrome C in *A. fumigatus* allowed long term persistence in the murine lung in an invasive pulmonary aspergillosis coupled with *in vitro* resistance to killing by a murine macrophage like cell line (18). Consequently, in this study, we utilized simultaneous tracking of two independent cell death markers over time, endogenous H2A:mRFP fluorescence and viability staining by Sytox Blue dead cell stain, to discover that the *A. fumigatus cycA* null mutant strain is also resistant to both chemically induced ROS *in vitro* and leukocyte induced ROS in a murine model of infection. Loss of *cycA* leads to increased conidial viability after treatment with H_2_O_2_ as measured by greater CFUs in a plating assay, retention of H2A:mRFP fluorescence, and reduction of Sytox Blue staining. Interestingly, we observe that in the CEA10 H2A:mRFP background strain, loss of H2A:mRFP fluorescence primarily occurs between 4hrs and 6hrs, while Sytox Blue staining occurs mainly between 6hrs and 8hrs. These results suggest an order of events during conidial cell death in which loss of nuclear integrity (measured as loss of mRFP fluorescence) precedes loss of cell viability during H_2_O_2_-induced conidial cell death.

Δ*cycA*-dependent conidial ROS resistance also has functional consequences during infection and leukocyte clearance. We observed that at 36 hpi, the Δ*cycA* strain displayed increased viability in both lung neutrophils and lung monocytes. Data from mice and humans identified an essential role for host ROS generated via NADPH oxidase for the clearance of *A. fumigatus* conidia from the lung (7, 8). Additionally, we observed that host ROS can induce pathways in *A. fumigatus* associated with cell death, and that the fungal anti-cell death protein, Bir1, counteracts this process (9). In a mixed bone marrow setting with p91phox^+/+^ and p91phox^−/−^ leukocytes in the same lung, we surprisingly observed a stark increase in viability in the Δ*cycA* strain in p91phox^-/-^ neutrophils compared to p91phox^+/+^ neutrophils. This data suggests that *cycA* not only confers resistance to host oxidative killing, but also to non-oxidative forms of conidial killing. The resistance of the Δ*cycA* strain to leukocyte killing may explain previous findings that it was able to persist in the murine lung up to 29 days post infection in immunocompromised models of infection despite a significant growth defect (18). The NADPH oxidase-independent factors contributing to conidial killing remain an area of active investigation.

It remains unclear whether there is a mechanistic relationship between Bir1 function and CycA in fungi. Our transcriptome analyses of the *A. fumigatus* Bir1 N-terminal domain over-expression strained revealed altered transcript levels of genes encoding proteins associated with the mitochondria and electron transport chain. For example, we observed changes in mRNA transcript abundance corresponding to genes critical for the incorporation of TCA cycle intermediates into Complex II of the electron transport chain. Increased mitochondrial respiration can lead to mitochondrial membrane leakage, increased endogenous ROS production, and an increased flow of electrons through ubiquinol-cytochrome b1c Complex III (43, 44). In this scenario, a possible explanation is that increased aconitate hydratase transcripts identified in our RNA-seq dataset are part of a compensatory increase in the activity of cytochrome C to maintain redox homeostasis. However, aconitate hydratase may also directly function in a protective role. This mitochondrial Fe-S cluster enzyme interconverts the TCA intermediates citrate, aconitate, and isocitrate. Increased aconitate hydratase activity results in increased reducing power to fuel oxidative respiration and increased flux through the TCA cycle that feeds into the electron transport chain at Complex II. Moreover, aconitate hydratase can also function as a ROS scavenger by converting the highly reactive intermediate *cis*-aconitate into the more stable product isocitrate, thereby reducing oxidative damage to cells (45). Further studies are needed to better understand the effect of aconitate hydratase on conidial cell death by H_2_O_2_ and leukocytes.

Thus, our data suggest a model whereby swollen *A. fumigatus* Bir1^OE-N^ conidia may exhibit increased survival upon H_2_O_2_ stress due to increased electron flow through the electron transport chain from Complex II to Complex III and Complex VI, increasing endogenous ROS production and subsequent detoxification enzymes (44). In this scenario, oxidative phosphorylation may continue from Complex III to Complex VI via electron transport from mitochondrial cytochrome b-c1 to cytochrome c, facilitating downstream ATP synthase activity. Previous work has shown cytochrome c activity as critical for ROS sensitivity (18), and taken together, the transcriptome of Bir1^OE-N^ cells and previous work centered around CycA both highlight the possibility that cytochrome c mediates overall metabolic adaptation in *A. fumigatus* and that loss of cytochrome c results in fungal conidia that are able to combat further oxidative stress exposure. This finding supports a hypothesis that altered Bir1 levels effect intrinsic ROS resistance through perturbation of proteins involved in mitochondrial function. However, we also cannot rule out the existence of a cell death effector in fungi that is regulated by CycA and Bir1, similar to the regulation of caspase activity by metazoan homologs of these 2 proteins.

Taken together, our data suggests that cytochrome c in *A. fumigatus* can mediate cell death induced by exogenous H_2_O_2_, as well as by leukocytes. While our data suggests that this resistance is a consequence of altered mitochondrial function, cytochrome c in metazoa functions as an initiator of caspase dependent cell death, and surviving (Bir1 in fungi) functions to negatively regulate these cell death proteases. Further investigation is required to determine if CycA, and also Bir1, directly participate in a regulated cell death specific pathway and if a mechanistic relationship between the two exists in which a cell death specific protease is involved. Understanding potential protein interactors of Bir1 and CycA during exposure to ROS may elucidate whether conidial cell death under these conditions is part of a regulated pathway or a consequence of an altered metabolic state. These areas of investigation would give insights into potential therapeutic targets for fungal infections and also into the origin of regulated cell death processes in pathogenic molds.

## Methods

### Strains and growth conditions

Mutant strains were generated in the CEA10/CBS144.89 strain (46). All strains were stored as conidia at -80°C in 25% glycerol. Culturing of strains was done using glucose minimal media (GMM). GMM media consists of 6 g/liter NaNO_3_, 0.52 g/liter KCl, 0.52 g/liter MgSO_4_·7H_2_O, 1.52 g/liter KH_2_PO_4_ monobasic, 2.2 mg/liter ZnSO_4_·7H_2_O, 1.1 mg/liter H_3_BO_3_, 0.5 mg/liter MnCl_2_·4H_2_O, 0.5 mg/liter FeSO_4_·7H_2_O, 0.16 mg/liter CoCl_2_·5H_2_O, 0.16 mg/liter CuSO_4_·5H_2_O, 0.11 mg/liter (NH_4_)_6_Mo_7_O_24_·4H_2_O, 5 mg/liter Na_4_EDTA, 10 g/liter glucose. pH of media is adjusted to 6.5. To make solid GMM media, 1.5% agar (Difco granulated agar, BD Biosciences) was added to the above-described recipe. GMM media was autoclaved prior to use. For flow cytometry assays, liquid GMM was prepared as stated above, autoclaved, and subsequently filter sterilized using a 0.2uM vacuum filter bottle system (Corning) to make optically clear GMM. To gather conidia, strains were cultured from -80°C stocks onto solid GMM and incubated for 3 days (4 days for the Δ*cycA* strain) at 37°C plus 5% CO2. Conidia were collected by addition of 0.01% Tween-80, followed by agitation with a cell scraper, and filtering through Miracloth.

### Culture conditions, RNA extraction, & RNA Sequencing of Bir1OE-N swollen conidia

Strains were plated onto LGMM and incubated for 72 hrs at 37°C, 5% CO_2_ in the dark. After collection using 0.001% Tween 80-H_2_O, conidia were enumerated using a hemacytometer. In technical triplicate assays, 1x10^6^ ATCC46645-H2A:mRFP and ATCC46645-H2A:mRFP Bir1OE-N conidia were inoculated in 100 mL of RPMI + 0.5 ug/mL Voriconazole (Sigma-Aldrich) and incubated at 50 RPM for 6 hrs at 37°C. Fungal cells were collected by centrifugation and were flash frozen in liquid N_2_. Samples were then processed for RNA extraction after bead beating for 1 min with 0.1 mm glass beads, followed by Trisure (Zymo Research) and chloroform extraction and purification through Qiagen columns (Qiagen RNAeasy Kit). RNA quantity and quality was estimated by Nanodrop and through gel electrophoresis before submission to the Dartmouth Genomics Core Facility. RNA was processed through Illumina NextSeq500 75 cycles after library preparation (Poly-A capture). Fasta files were further analyzed using FungiDB and EuPathDB Galaxy tools, mapping reads onto the closest related annotated genome, “A1163 Aspergillus fumigatus” (47). Resulting DeSeq2 from the comparison of “ATCC46645-H2A:mRFP” vs. “Bir1-N” data is appended in supplementary Table 1. Differential expressed genes are visualized in Figure 1b.

### Mutant Strain Generation

CEA10:H2A.X*^A. nidulans^*∷*ptrA* (CEA10 H2A:mRFP) was constructed in two steps. First, *gpdA*(p) was amplified from DNA isolated from strain *A*. *nidulans* A4 (Source:FGSC) using primers RAC2888 and RAC2799. Histone variant H2A.X (AN3468) was amplified (without stop codon) from A4 DNA (primers RAC4582 and RAC4583). mRFP fragment was amplified from plasmid pXDRFP4 (Source:FGSC) (primers RAC2600 and RAC4575) and terminator for *A*. *nidulans trpC* gene was amplified (primers RAC2536 and RAC2537). The four fragments were then fused together via PCR (primers RAC1981 and RAC4134), resulting in H2A.X first round fragment as described in (48). Secondly, we targeted integration of the H2A.X∷rfp to the intergenic locus between AFUB_023460 and AFUB_023470 on chromosome 2. For this, a left homology arm (primers RAC3873 and RAC3874) and right homology arm (primers RAC3875 and RAC3876) was amplified from CEA10 genomic DNA. Dominant selection marker gene *ptrA* conferring resistance to pyrithiamine hydrobromide was amplified from plasmid pSD51.1 (primers RAC2055 and RAC2056). The four fragments were then fused together via PCR (primers RAC3877 and RAC3878) as described earlier (48). After the construct generation, polyethylene glycol mediated transformation of protoplast was performed as described earlier (48). mRFP fluorescence was confirmed with FACS (fluorescence activated cell sorting) analysis. To generate CEA10 H2A:mRFP Δ*cycA* strain, PCR was used to amplify approximately 1kb upstream (primers RAC5242 and RAC5243) and 1kb downstream (primers RAC5244 and RAC5245) of the *cycA* gene. These upstream and downstream elements were fused to the hygromycin resistance gene (HygR) via PCR (primers RAC5246 and RAC 5247) to generate a knockout construct. This knockout construct was transformed into CEA10 H2A:mRFP protoplasts. Protoplasts were plated onto 15mL of SMM (GMM plus 1.2M sorbitol) in 5mL of SMM top agar (SMM + 0.7% agar). The following day a 5mL layer of SMM top agar supplemented with hygromycin B (VWR) was added for a final concentration of 175ug/mL hygromycin for the entire 25mL SMM plate. To generate the *cycA* reconstituted strain (*cycA*+), the *cycA* loci including ∼1.3kb upstream and 200bp downstream of the open reading frame was amplified out of the genome by PCR (RAC5246 and RAC5419). As the Δ*cycA* strain displays nearly no growth on media with glycerol as the only carbon source (18), this amplified PCR product was used as the full reconstitution construct and transformants were selected for on SMM agar with 1% glycerol instead of 1% glucose as the carbon source. For all transformations, protoplasts were generated treating germlings derived from liquid cultures grown at 28°C for 11hrs with lysing enzyme from *Trichoderma harzianum* (Sigma). Transformants were screened by PCR and confirmed with southern blot analysis.

### *cycA* Sequence Annotation Correction and Multiple Sequence Alignment

The *cycA* nucleotide and amino acid sequences for the reference strain AF293 (Afu2g13110) and CEA10 (AFUB_028740) were downloaded from fungiDB (47). Using the AF293 nucleotide sequence, we identified introns and a new stop codon based on reads identified in the Kowalski et al. 2019 RNA seq data set (23). These newly determined introns were then verified with the combined AF293 RNA seq plot found on fungiDB. We then compared the nucleotide sequences of the updated AF293 introns and exons to that of CEA10 to determine any difference in amino acid sequences between the two. Protein structure modeling was done with Phyre2 (49)by comparing the updated AF293 *CycA* sequence to that of the human *CycS*. Protein sequences for orthologs of Cytochrome c were obtained from UniProt or FungiDB and analyzed for similarity by multiple sequence alignment using ClustalW 2.1 and phylogeny.fr (50). Alignment was edited for clarity using JalViewJS (51).

### Cell Death Assays

2 x 10^6^ conidia/mL were swollen at 37°C for 4 hrs in 10mL cultures using optically clear LGMM. As the loss of *cycA* results in swelling defect, Δ*cycA* strains were swollen for an additional 2 hrs (Supplementary Figure 2). After swelling, cultures were vortexed to resuspend and 1mL of swollen culture was reserved for each strain as an untreated (0hr) control. For 2 hr, 4 hr, 6 hr, and 8 hr H_2_O_2_ – treatment cultures, 1 mL of swollen conidia was aliquoted into a 5 mL screwcap centrifuge tube (VWR). To each 1 mL aliquot, 1 mL of 20mM H_2_O_2_ in optically clear LGMM was added, for a final concentration of 10^6^ conidia/mL and 10 mM H_2_O_2_ in 2mL. H_2_O_2_ – treated cultures were incubated in 37oC, and every 2 hrs a 2 mL treatment culture was stained with Sytox blue dead cell stain (Invitrogen) at a concentration of 1 uM per manufacturer’s protocol and incubated at room temperature for 5 min. After staining, conidia were analyzed by flow cytometry. For flow cytometric analysis, 15,000 events were gathered for each strain at each timepoint. To process flow cytometry data, events were visualized on FSC-A vs FSC-H plots and gated for single events. Gating of events for mRFP florescence was determined by using conidia lacking mRFP as a negative control for gating and untreated (0hr) H2A:mRFP conidia as a positive control. For Sytox blue staining, a negative control for gating was used in which untreated conidia were stained with Sytox blue dead cell stain. Based on these gates, the percent of mRFP positive and Sytox blue positive singlets were derived for each treatment timepoint and compared to untreated (0hr) florescence. All flow cytometry was conducted on a Cytoflex S cytometer (Beckman Coulter), and flow cytometry data analysis was done on FlowJo version 10.8.1.

### CFU Assay

2 x 10^6^ conidia/mL were swollen at 37°C for 4 hrs in optically clear LGMM. After swelling, conidia were treated with 5 mM H_2_O_2_ for 2.5 hrs. After treatment, conidia were centrifuged to remove H_2_O_2_ media and resuspended in 0.01% Tween 80 in water. Conidia were enumerated using a hemacytometer and 100 conidia were plated in 5 mL of GMM top agar (GMM with 0.7% agar) overlaid (?) on 20 mL of GMM agar. Cultures were incubated at 37°C for 36 hrs and colonies were counted. The number of colonies from H_2_O_2_ – treated cultures was then compared to the strain specific untreated control groups.

### Mice

C57BL/6J mice (strain #: 000664) and p91phox^-/-^ mice (strain #: 002365) were purchased from The Jackson Laboratory. C57BL/6.SJL (WT CD45.1^+^) were purchased from Charles River Laboratories (strain # 00564) and crossed with C57BL/6 (CD45.2^+^) to generate CD45.1^+^CD45.2^+^ recipient mice for bone marrow chimeric mice.

### In Vivo FLARE

To generate FLARE conidia, 7.5 x 10^8^ CEA10:H2A, *ΔcycA*, and *ΔcycA+* conidia were rotated in 0.5 µg/ml Sulfo-NHS-LC-Biotin (A39257; ThermoScientific) in 1 ml 50 mM carbonate buffer (pH 8.3) for 1 hr at RT, washed in 0.1 M Tris-HCl (pH 8), labeled with 0.02 mg/ml Streptavidin, Alexa Fluor 633 conjugate (S-21375; Molecular Probes) in 1 ml PBS for 30 min at RT, and resuspended in PBS and 0.025% Tween 20 for experimental use. For intratracheal challenge, mice were lightly anesthetized with isoflurane and immobilized in an upright position using rubber bands attached to a Plexiglas stand. The indicated number of conidia (typically 3x10^7^) in a volume of 0.05 ml PBS, 0.025% Tween 20 were delivered to the lung using a micropipette. Single cell lung suspensions were prepared for flow cytometry as previously described^1^, with minor modifications. Briefly, perfused murine lungs were placed in a gentle MACS C tube and mechanically homogenized in 5 ml of PBS using a gentle MACS Octo Dissociator (Miltenyi Biotecc). Lung cell suspensions were lysed of RBCs, enumerated, and stained with fluorophore-conjugated antibodies prior to flow cytometric analysis on a Beckman Coulter Cytoflex LX and analyzed with FlowJo version 10.8.1. Neutrophils were identified as CD45^+^ CD11b^+^ Ly6G^+^ cells, inflammatory monocytes as CD45^+^ CD11b^+^ CD11c^-^ Ly6G^-^ Ly6C^hi^ cells, and Mo-DCs as CD45^+^ CD11b^+^ CD11c^+^ Ly6G^-^ Ly6C^hi^ MHC class II^+^ cells. As previously described^2^, phagocytes that contain live conidia are RFP^+^ and AF633+ (R1) and phagocytes that contain dead conidia are RFP^-^ AF633^+^ (R2). Conidial phagocytosis was quantified as the sum of the fraction of a given phagocyte in the R1 gate and the fraction of a given phagocyte in the R2 gate (R1+R2). To assess how effective phagocytes were at killing conidia, the fraction of viable conidia was calculated as R1/(R1+R2).

### Mixed bone marrow chimera

Lethally irradiated (900 cGy) F1 progeny (from a cross of C57BL/6 and C57BL/6.SJL strains) were reconstituted with a mixture of 1-2.5 x 10^6^ CD45.1^+^ p91phox^+/+^ and 1-2.5 x 10^6^ CD45.2^+^ p91phox^-/-^ bone marrow cells, treated with enrofloxacin in the drinking water for 21 days to prevent bacterial infections, and rested for 6 weeks prior to use. Mice were challenged with 3x10^7^ WT, Δ*cycA*, or Δ*cycA*+ FLARE conidia and after 36 hours were analyzed by flow cytometry as described above.

### Data Availability

Raw and processed RNA sequencing data in this study are available at the GEO repository under accession number GSE233942.

All diagrams were generated using BioRender.com. All data was processed using Prism GraphPad version 9.5.1.

## Acknowledgements

Dartmouth Genomics Core Facility (NCI Cancer Center, Support Grant 5P30CA023108-37, RRID:SCR_021293. Flow cytometry experiments were carried out in DartLab, the Immune Monitoring and Flow Cytometry Shared Resource at the Norris Cotton Cancer Center at Dartmouth, with NCI Cancer Center Support Grant 5P30 CA023108-41 and at Memorial Sloan Kettering Cancer Center, with NCI Cancer Support Grant P30 CA008748. This work was supported by NIH R01 AI139632 (to TMH and RAC), NIH R37 AI093808 (TMH), and NIH F31 AI 161996 (MAA).

Thanks to Dr. Charles Puerner for graphics assistance and Dr. Joshua Kerkaert for assistance on intron prediction and annotation correction of *cycA*.

---

**Supplementary Figure 1.** Gating strategy for lung FLARE experiments. (A) Gating strategy to identify lung phagocyte subsets at 36 hpi. (B) The plots indicate the number of lung phagocytes at 36 hpi from mice of the indicated genotypes. Data are representative of 2 independent experiments. (B and D) Dots represent individual mice and data are expressed as mean ± SEM. Statistics: (B) Kruskal-Wallis test with Dunn’s multiple comparisons test.

**Supplementary Figure 2.** Conidial size of CEA10 H2A:mRFP and Δ*cycA* strains as determined by flow cytometry. The Δ*cycA* strain displays similar size on forward/side (FSC/SSC) gating at 6hrs to the CEA10 H2A:mRFP strain at 8hrs.

